# From DNA Helicity to Structural Twist: An Undergraduate Laboratory Module for DNA Origami Design

**DOI:** 10.64898/2026.11.18.767739

**Authors:** Seongmin Seo, Anirudh S. Madhvacharyula, Sihyun Kim, Jong Hyun Choi

## Abstract

Precise control of nanoscale architectures makes DNA origami a promising platform for biomedical applications such as drug delivery, biosensing, and bioimaging. A key constraint in DNA origami design is that crossover placement must account for the helical pitch of DNA. For new learners, however, it can be difficult to understand how a DNA helical pitch affects assembled structures. Here, we report a hands-on laboratory module to teach undergraduate students about this fundamental helical pitch/structure relationship. A rectangular DNA origami tile is designed as a model system with its helical pitch varying from 10.5 to 10.84 base-pairs per turn. When the tiles are linked into a ribbon, the small mismatch-driven deformations of individual tiles accumulate into a global twist. The twist appears as characteristic kinks under atomic force microscopy (AFM). In a five-day pilot, three engineering students designed the DNA origami tile and used CanDo finite-element simulations to predict its deformations at different helical pitches. They assembled ribbons and used an intercalator, ethidium bromide (EtBr), to vary DNA helicity without changing the crossover layout. With increasing EtBr concentration, all three participants observed ribbons’ chirality transition from right-handed to flat to left-handed conformations. By comparing predictions with their own observations, students gain a practical understanding of how DNA helicity governs DNA structure design.

## INTRODUCTION

DNA nanotechnology exploits sequence-specific base pairing to assemble DNA molecules into predesigned structures with nanometer-scale precision^1, 2^. Building on this ability, researchers have designed various DNA nanostructures for biomedical applications. At its simplest, architected DNA can arrange recognition elements at defined locations for biosensing^3, 4^. Fluorescent probes extend this capability to cells, selectively labeling cancer cells^5^, resolving multiple subcellular structures in one sample^6^, and tracking individual endocytic carriers^7^. The structure itself can also contribute to biological function. DNA nanocages protect enclosed enzymes from protease digestion and can enhance their catalytic activity^8^. Rectangular DNA constructs preferentially accumulated in the kidneys and showed protective effects in a mouse model of acute kidney injury^9^. Beyond fixed geometries, reconfigurable DNA nanostructures can change shapes to perform specific tasks in response to external cues. Barrel-shaped nanorobots reveal their payload upon recognizing a molecular target^10^. Another dynamic device delivered thrombin to tumor vasculature in vivo^11^. A deployable transformer expands to create nanopores that regulate payload release through lipid membranes^12^. In each of these examples, function follows geometry that is set at the design stage.

DNA origami is a widely used method for building such structures^13, 14^. In this approach, numerous short staple strands fold a long, single-stranded scaffold into complex shapes. Each staple binds distant segments of the scaffold and brings them together forming crossovers, where a DNA strand passes from one helix to the next. The placement of these crossovers is a central consideration in origami design. As the DNA backbone rotates along the helix, the helical pitch determines how the crossover sites line up. If the crossover spacing is not compatible with the helical pitch of natural B-form DNA, the mismatch can generate internal stress and cause the assembled structure to bend or twist away from its intended geometry^15, 16^.

To facilitate the design of complex structures, computer-aided design tools such as caDNAno^17^ and scadnano^18^, along with automated platforms including PERDIX^19^, ATHENA^20^, and MagicDNA^21^, have been developed. Computational tools, including finite-element models (CanDo^22^ and SNUPI^23^) and molecular dynamics (MD) models (oxDNA^24^ and MrDNA^25^), predict structural conformations before synthesis. Although these tools have made DNA nanostructure design more accessible, they can obscure the underlying design principles for new learners. Design software places crossovers at positions consistent with an assumed helical pitch, allowing users to create valid designs without understanding why those positions depend on DNA helicity or how a helicity mismatch leads to global structural deformation. Observing such deformation experimentally is also challenging. Most structural features of DNA nanostructures lie below the optical diffraction limit. Atomic force microscopy (AFM) can resolve the issue, but images of surface-bound structures do not capture their 3D shape in solution.

Here, we present a hands-on laboratory module that helps students see how this molecular design constraint affects the shape of DNA origami assemblies. Students explore this relationship through a full research workflow that includes design and finite-element simulation with experimental assembly and AFM characterization. The activities build on our earlier study of DNA origami tiles^26, 27^ and ribbons^28^. When DNA origami tiles are linked into a ribbon, small deformations in individual tiles caused by helicity mismatch accumulate into a global twist of the ribbon. This twist appears as a characteristic pattern of kinks in AFM images. By varying the concentration of the intercalator ethidium bromide (EtBr), students can observe how changes in DNA helicity alter the ribbon shape without changing the crossover layout.

Previous laboratory modules have made different aspects of DNA nanotechnology accessible to students. Beshay et al. used a previously designed DNA origami nanorod to teach folding and gel-based characterization^29^. Other undergraduate exercises use gel electrophoresis to examine topology changes in DNA nanoswitches^30^ or quantify strand displacement in switchback DNA^31^. More recently, Talbot et al. used isothermal assembly and gel electrophoresis to teach students how to quantify the assembly yield of double-crossover motifs^32^. Our module extends these laboratory exercises by integrating design and simulation with experimentation. Students examine an origami design, use simulations to predict structure and deformation, and compare their predictions with their own AFM observations. The module highlights a design principle not addressed in the exercises described above, that is, DNA helicity constrains crossover placement, and the constraint can shape the assembled structure.

This five-day instructional laboratory module was designed for upper-level undergraduates in their junior year or later. We piloted it with three students who had recently earned undergraduate degrees in engineering. Students work with tile designs in caDNAno and use CanDo to predict how the tiles deform at different helical pitches (Fig. 1a). They then assemble the tiles into ribbons, treat them with EtBr, and examine the geometry by AFM. By linking molecular geometry to observable changes in structure, this module gives students a foundation for reasoning about DNA nanostructure design and evaluating design choices through simulation and experiment.

**Figure 1.**
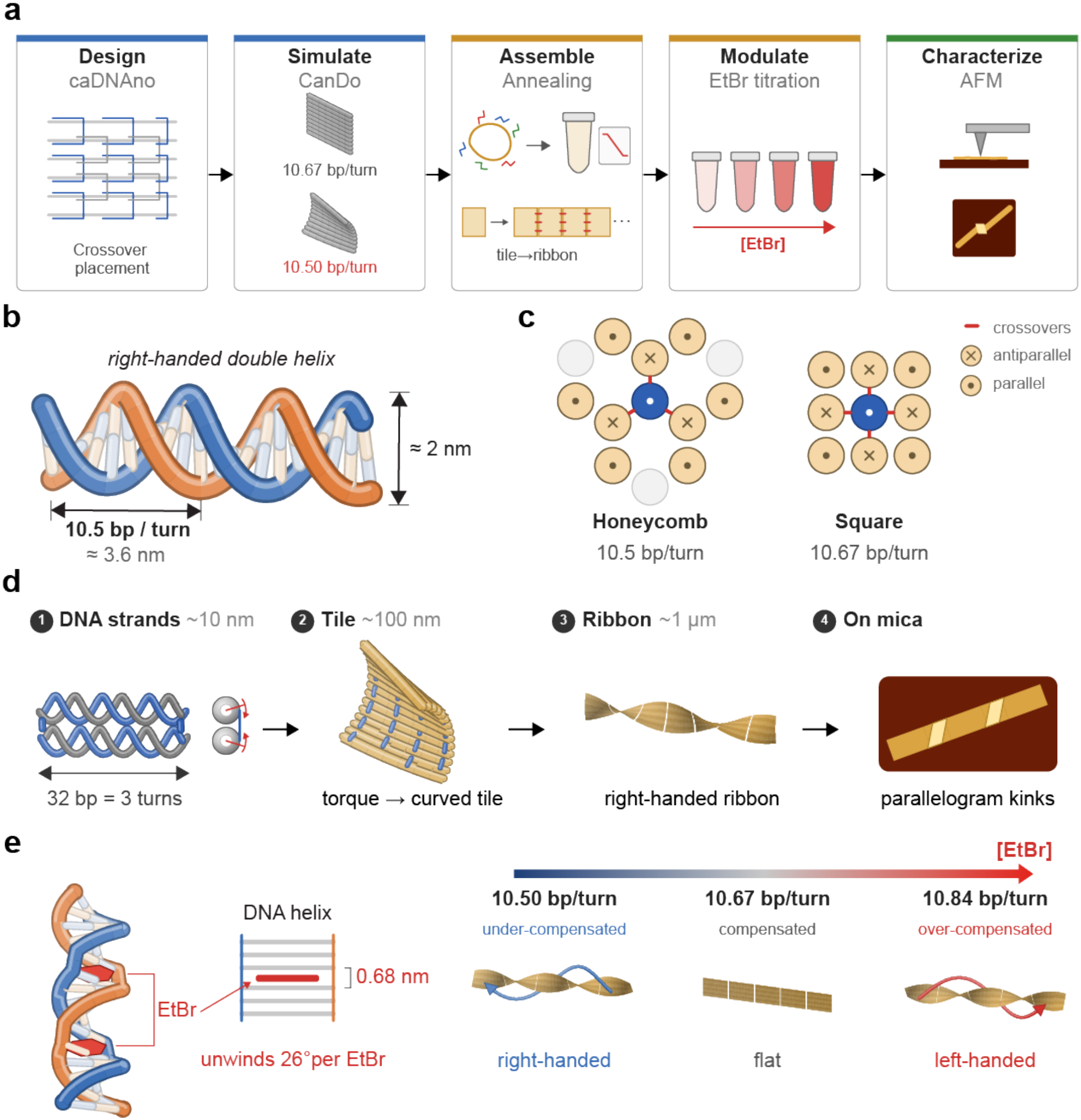
Educational workflow and conceptual framework of DNA helicity and origami ribbon chirality. **a**, Laboratory workflow integrating DNA origami design, CanDo simulation, assembly, EtBr-mediated modulation, and structural characterization. **b**, Geometry and helical pitch of Watson-Crick B-form DNA. **c**, Crossover arrangements in honeycomb and square lattices for DNA origami design. **d**, Helicity mismatch induces local deformation in DNA origami tiles, resulting in global ribbon twisting and characteristic kinks. **e**, EtBr intercalation modulates DNA helicity and ribbon conformation through under-compensation, compensation, and over-compensation of the mismatch.

## CONCEPTUAL BACKGROUND

### DNA Helicity and Origami Design

Watson-Crick B-form DNA is a right-handed double helix with a diameter of approximately 2 nm and a helical pitch of approximately 10.5 base pairs per turn (bp/turn) as illustrated in Fig. 1b. In DNA origami, multiple DNA duplexes are connected by crossovers^33, 34^ to form structural elements such as edges and sheets. Since the DNA backbone rotates along the helical axis, crossover positions must align with the orientation of neighboring helices. Two commonly used duplex arrangements are the square and honeycomb lattices (Fig. 1c). Each lattice uses a specific DNA helical pitch to set the spacing between allowable crossover sites.

A honeycomb lattice has up to three neighbors arranged at 120° intervals^14^. It assumes two-thirds of a turn every 7 bp, corresponding to 10.5 bp/turn. Crossovers to the same neighbor repeat every 21 bp (two turns). The honeycomb lattice therefore matches the natural helicity of B-form DNA. In a square lattice, each helix has up to four neighbors arranged at 90° intervals^35^. The design assumes three-quarters of a helical turn every 8 bp, corresponding to 10.67 bp/turn. Crossovers to the same neighbor repeat every 32 bp (three turns). As a result, the square lattice introduces a small mismatch between the designed and relaxed helical pitches.

This mismatch can be quantified by the mismatch angle α, defined as the rotational difference between a relaxed 32-bp DNA segment and the three turns required by the design^28^. For a 32-bp segment in a square-lattice design,

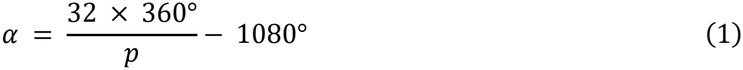

where *p* is the helical pitch in bp/turn. For relaxed B-form DNA, this angle is α ≈ +17°. This mismatch is small at each segment but recurs throughout the structure. Forcing all domains in a 32-helix, single-layer rectangular tile to adopt the designed pitch would cost ∼32 kcal/mol of elastic energy^26, 27^. Instead, the tile bends and twists out of plane to relieve strain.

### From DNA Helicity to Ribbon Chirality

Since a DNA origami tile flattens on a mica surface, it is difficult to resolve its 3D deformation state in free solution by AFM. The deformation can be amplified by connecting the tiles with linker strands to form an elongated ribbon (Fig. 1d). The local deformation of each tile then accumulates along the ribbon, where twisting is coupled with bending in the single-layer ribbon, producing a spiral conformation^28^.

The handedness of this global twist is referred to as *ribbon chirality*. For the square-lattice design the natural helicity of B-form DNA (*α* > 0) produces a right-handed ribbon. Reversing the sign of *α* would produce a left-handed ribbon. Eliminating the mismatch (*α* = 0) should result in a flat ribbon. When a twisted ribbon is deposited onto mica for AFM imaging, parts of the ribbon flatten against the surface. The remaining twist becomes concentrated into double-layer, parallelogram-shaped kinks. Right- and left-handed ribbons form mirror-image kink patterns, allowing their handedness to be distinguished from AFM images. This relationship was previously verified through biotin–streptavidin labeling and supported by CanDo simulations^28^.

### Intercalator-Mediated Modulation of DNA Helicity

DNA helicity can be adjusted after origami assembly using DNA intercalators. These molecules insert between adjacent base pairs and unwind the double helix, increasing its helical pitch in bp/turn. EtBr is a commonly used intercalator that unwinds DNA by approximately 26° per bound molecule^36, 37^. The extent of unwinding depends on the intercalation density (*ρ*), defined as the number of bound molecules per base pair^28^. For the 32-bp crossover design, compensating the helicity mismatch of ∼17° requires an estimated density of *ρ* ≈ (17°/32)/26° ≈ 1/48 bp^−1^. The EtBr concentration required to reach this density is determined by the binding equilibrium, which depends on the DNA concentration. Below ∼0.1 molecule/bp, *ρ* increases approximately linearly with EtBr concentration. On this basis, compensation was predicted at ∼0.75 µM EtBr for 2 nM ribbons, close to the ∼1 µM at which planar ribbons without any kinks were observed^28^.

As EtBr binding increases, a ribbon progresses through three regimes: under-compensation, compensation, and over-compensation (Fig. 1e). The right-handed twist decreases until the ribbon becomes approximately flat, and further intercalation reverses the twist to left-handed. This reconfiguration is complete within approximately 10 minutes. This process can be reversed by adding excess DNA duplexes, which can compete for bound EtBr and restore the original ribbon conformation^28^.

Similar changes have been demonstrated using other intercalators, including TMPyP4 and chloroquine^28^. In contrast, the minor-groove binder DAPI increases ribbon twisting and can induce tubular conformations. DNA origami geometry may also be modified through base-pair insertions or deletions between crossovers during design^15^. In this module, we use EtBr to progressively modulate DNA helicity and compare the resulting ribbon geometries with CanDo simulations at different helical pitches.

## RESULTS

### DNA Origami Tile Design and Finite Element Simulation

To investigate how DNA helicity affects structural geometry, students designed a single-layer rectangular DNA origami tile consisting of 32 parallel double helices using caDNAno (Fig. 2a). The square-lattice design has crossovers spaced every 32 bp, corresponding to ∼10.67 bp/turn. Single-stranded domains were placed at both lateral edges for subsequent ribbon assembly. Because relaxed B-form DNA does not match this designed pitch, every 32-bp segment carries a helicity mismatch (Fig. 2b).

**Figure 2.**
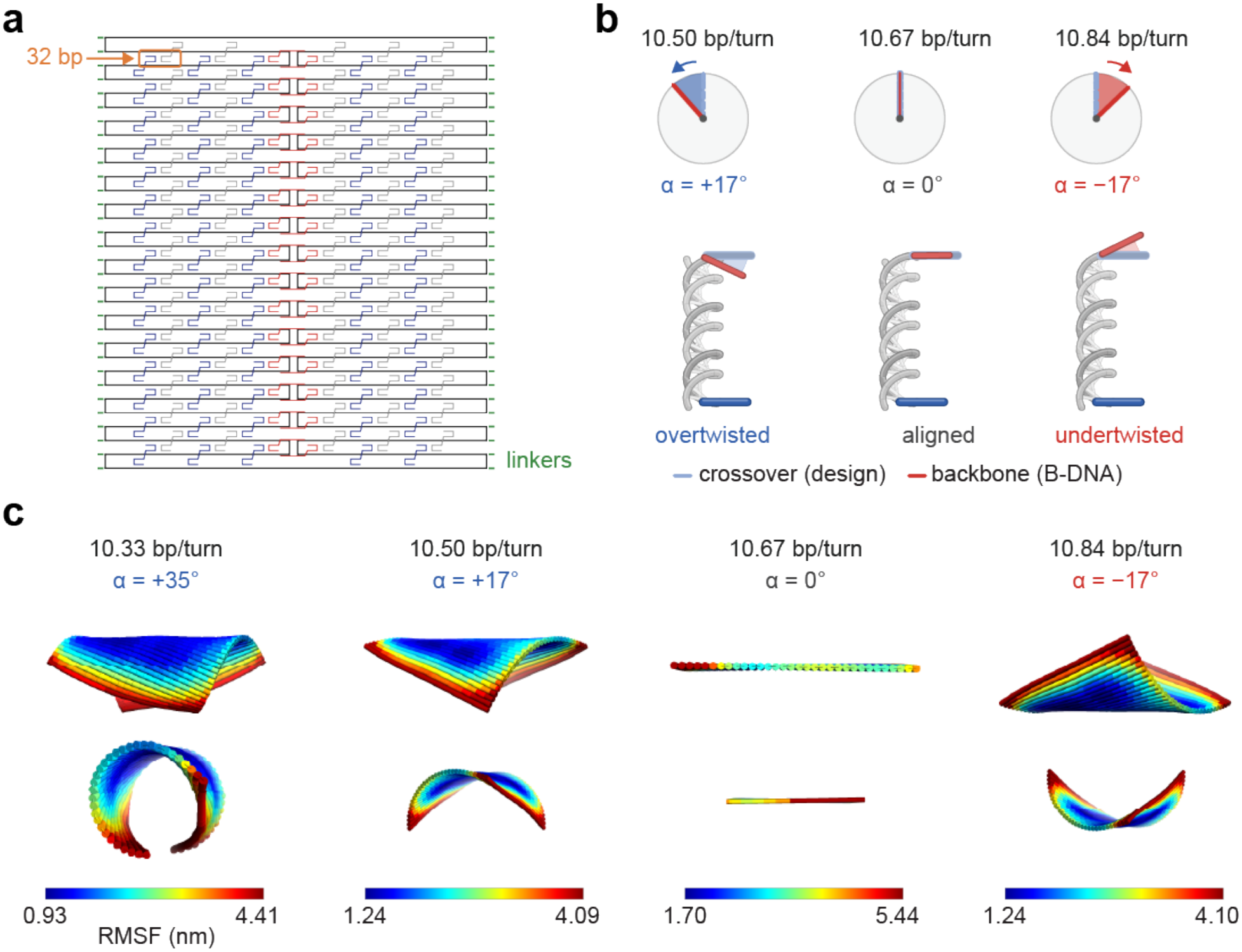
Helicity-dependent structural deformation. **a**, caDNAno design of the rectangular DNA origami tile with crossovers spaced every 32 bp. **b**, Schematic illustration of helicity mismatch between the designed (10.67 bp/turn) and relaxed DNA helical pitches. **c**, CanDo finite-element simulations of individual DNA tiles at different helical pitches (10.33, 10.50, 10.67, and 10.84 bp/turn). Color maps indicate the root-mean-square fluctuation (RMSF) of each structure in nanometers.

Using CanDo finite-element modeling^22^, students simulated the tile at four helical pitches: 10.33, 10.50, 10.67, and 10.84 bp/turn. These correspond to *α* = +35°, +17°, 0°, and −17°, respectively (Fig. 2c). The crossover arrangement remained unchanged across all simulations. At the designed pitch of 10.67 bp/turn, the tile maintained an approximately flat conformation. At 10.50 bp/turn, corresponding to natural B-form DNA, the tile bent and twisted out of plane. Further decreasing the pitch to 10.33 bp/turn increased the curvature, producing a nearly tubular conformation. In contrast, a pitch of 10.84 bp/turn reversed the direction of bending, with an extent of deformation comparable to that at 10.50 bp/turn. The color maps show the root-mean-square fluctuations (RMSF), representing local flexibility rather than the extent of deformation. These results demonstrate how the magnitude and direction of helicity mismatch affect tile geometry and provide a basis for interpreting ribbon chirality in subsequent experiments.

### Assembly and Characterization of DNA Tiles and Ribbons

Students assembled two rectangular DNA origami tiles (A and B) through thermal annealing (Fig. 3a; see Methods). Both tiles share the same scaffold routing and staple arrangement but have distinct DNA sequences. To polymerize the tiles, sequence-specific linker strands were used to connect two separate tiles. Each linker set binds designated edges of adjacent tiles, allowing stepwise assembly into dimers and elongated ribbons.

**Figure 3.**
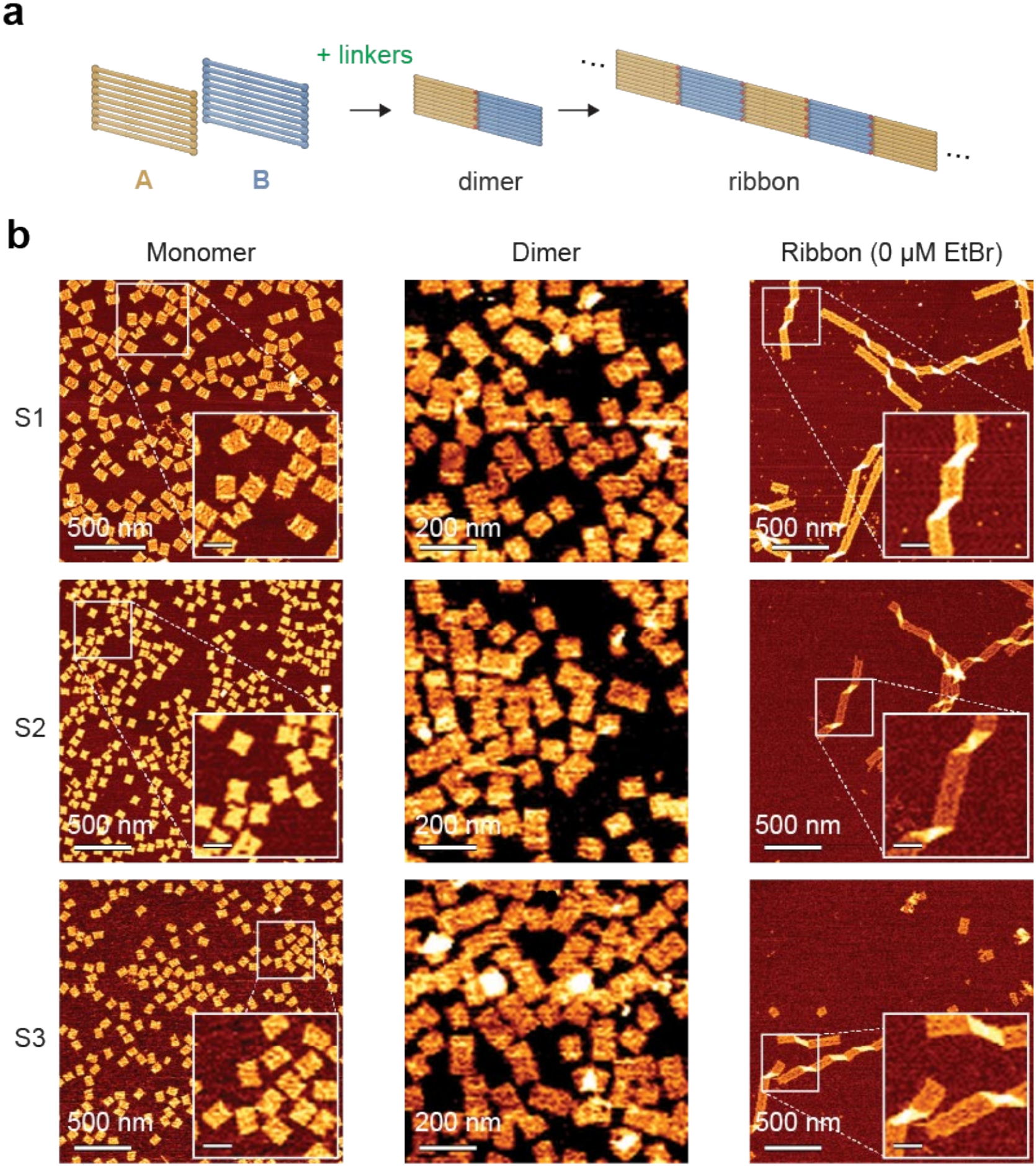
Assembly and characterization of DNA origami tiles and ribbons. **a**, Schematic illustration of the hierarchical assembly of DNA origami tiles into dimers and elongated ribbons using linker strands. **b**, AFM images of monomers, dimers, and ribbons obtained by three students (S1, S2, and S3). Insets show magnified views of representative structures. Scale bars in insets denote 100 nm.

All three students (S1, S2, and S3) characterized their assembled structures using AFM as shown in Fig. 3b. The monomer samples showed well-formed rectangular tiles, while the dimer samples contained pairs of connected tiles. Further assembly produced elongated ribbons, some extending to micrometer-scale lengths. In the absence of EtBr, the ribbons exhibited parallelogram-shaped kinks, as highlighted in the AFM images (Fig. 3b, insets).

### Helicity-Dependent Structural Conformation

To examine the effect of DNA helicity on ribbon geometry, participants treated the assembled ribbons with different concentrations of EtBr. Figure 4a illustrates the expected transition from right-handed to flat and left-handed ribbon conformations. Each participant characterized the ribbons using AFM at EtBr concentrations of 0, 0.25, 1, and 2 μM (Fig. 4b). Without EtBr, the ribbons exhibited dense right-handed kinks. At 0.25 μM, the kinks became less frequent but retained their handedness. At 1 μM, the kinks disappeared mostly, indicating a flat conformation. Further increasing EtBr to 2 μM produced mirror-image kinks associated with left-handed twisting. All three participants observed similar changes in ribbon morphology (Fig. 4b).

**Figure 4.**
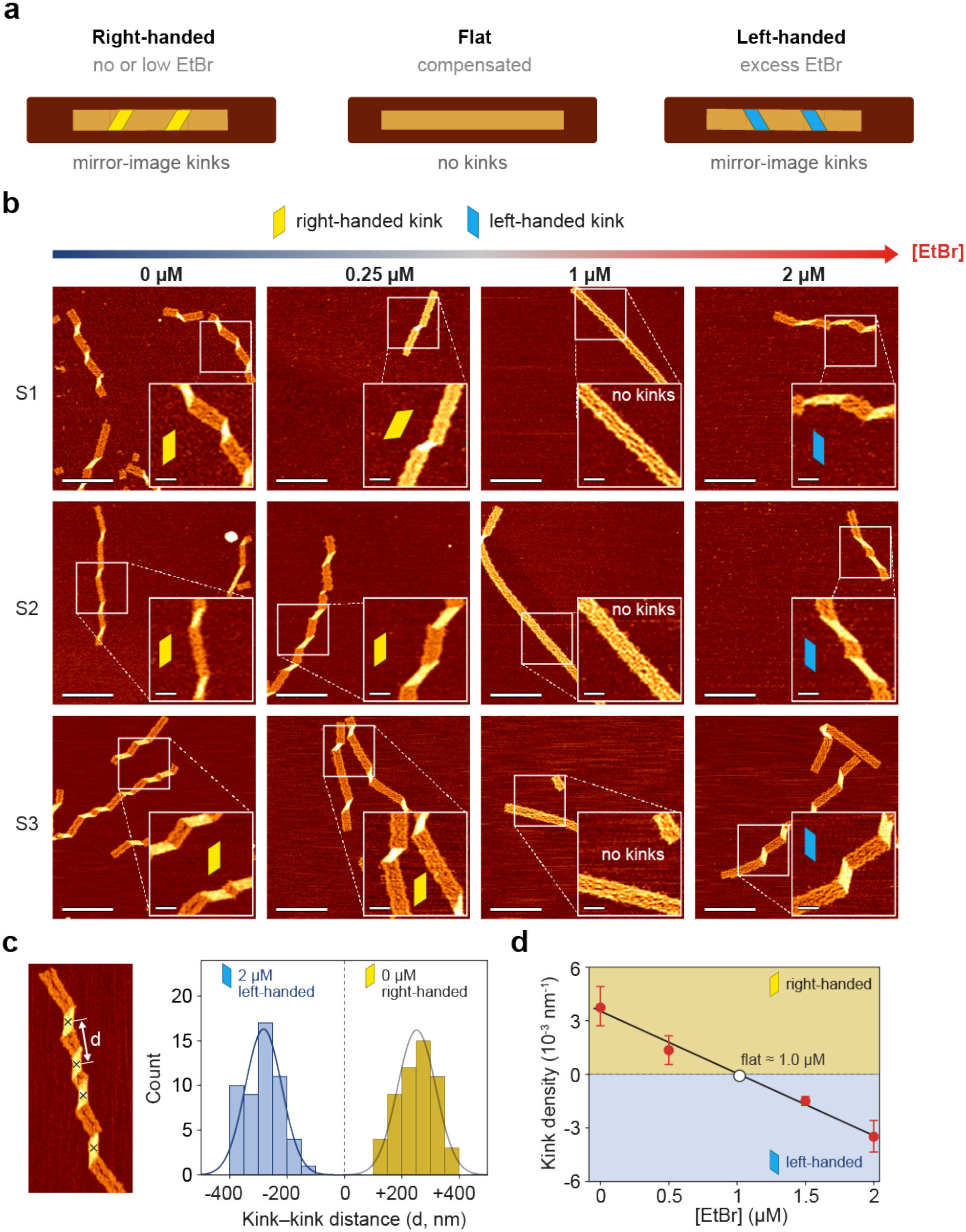
EtBr-induced modulation of DNA ribbon chirality. **a**, Schematic illustration of right-handed, flat, and left-handed ribbon conformations and their corresponding AFM kink morphologies. **b**, AFM images of DNA ribbons prepared by three participants (S1–S3) at various EtBr concentrations (0, 0.25, 1, and 2 μM). The images indicate the disappearance of right-handed kinks and the emergence of left-handed kinks with increasing EtBr concentration. Yellow and blue markers denote right-handed and left-handed kinks, respectively. Scale bars are 500 nm in the main images and 100 nm in the insets. **c**, Distributions of signed kink-to-kink distances at 0 and 2 μM EtBr, replotted from previously reported data^28^. Here n denotes the number of measured distances between adjacent kink centers (n = 54 at 0 μM and n = 52 at 2 μM). Solid curves represent Gaussian fits. **d**, Kink density as a function of EtBr concentration^28^. Points represent the inverse of the mean kink-to-kink distance, and error bars are the standard deviation. The open circle marks the zero crossing of the linear fit (∼1.0 μM), where the ribbons become flat.

To demonstrate how ribbon twisting can be quantified, we used kink-distance data previously reported^28^ for the same ribbon system as an instructional example (Fig. 4c,d). In those data, distances (d) between adjacent kink centers were assigned positive and negative values for right- and left-handed kinks, respectively. The distributions at 0 and 2 μM EtBr were approximately Gaussian, centered near +250 and −280 nm, respectively (Fig. 4c). Kink density, calculated as the inverse of the mean signed kink-to-kink distance, decreased approximately linearly with EtBr concentration, from approximately ∼3.8 × 10^−3^ nm^−1^ at 0 M to −3.6 × 10^−3^ nm^−1^ at 2 μM (Fig. 4d). It passes zero near 1 μM, consistent with the flattening observed by the participants. This reversal was qualitatively consistent with the opposite tile deformations predicted by CanDo simulations in Fig. 2c. We used this example to show participants how AFM observations can be quantified to characterize ribbon twisting.

## DISCUSSION

### Participant Learning Outcomes

All three participants completed the laboratory activities and generated their own datasets on DNA tiles, ribbons, and EtBr-treated ribbons (Figs. 3b and 4b). Instructor observations indicated that participants could identify ribbon handedness from AFM images and relate the changes to helicity mismatch. By comparing CanDo predictions with experimental observations, students analyzed the results and drew conclusions.

Participant progress reports provided further evidence of their conceptual reasoning. One participant (S1) independently extended the CanDo simulations with different design by varying the helical pitch from 10.3 to 10.7 bp/turn. The student reported stronger right-handed deformation at lower pitches and a shift toward left-handed deformation at 10.7 bp/turn. This participant also calculated the intrinsic DNA twist and crossover angle for each condition. The analysis demonstrated an effort to connect molecular helicity with tile geometry.

Another participant (S2) examined the effects of mechanical parameters on the predicted deformation. Increasing torsional stiffness in CanDo simulation produced greater tile twisting, whereas changing bending stiffness had little effect. The student attributed the latter observation to the absence of external forces. However, this explanation did not account for the internal stress generated by helicity mismatch, highlighting an area requiring further instruction. Overall, the observations and progress reports provide preliminary evidence of participants’ understanding of DNA helicity, crossover placement, and structural deformation toward DNA origami design.

### Educational Implications

The module addresses a gap left by design automation: connecting molecular-scale DNA helicity to the geometry of assembled structures. It uses ribbon chirality as an observable indicator of helicity-dependent deformation. Students relate subtle changes in DNA helicity to distinct (right-handed, flat, or left-handed) ribbon conformations and can extend this qualitative observation to quantitative analysis using kink density (Fig. 4c,d). Students can apply this understanding when selecting lattice geometries, adjusting crossover spacing, or evaluating structural deformation through simulations and experiments. The same principles govern dynamic DNA nanostructures^12, 38^, whose conformations can be controlled by internal strain or DNA-binding molecules.

Two design choices made the module accessible to participants with limited experience in DNA nanotechnology. First, we used a single-layer rectangular tile with a repeating square-lattice layout. Participants can identify the crossover pattern in caDNAno and perform simulations early in the five-day module. The repeating 32-bp crossover spacing also makes it easy to trace how a local helicity mismatch deforms individual tiles and how those deformations accumulate in a ribbon twist.

Second, we used finite-element modeling because the exercise focuses on equilibrium shape. Coarse-grained MD tools such as oxDNA can capture finer structural detail and time-dependent behavior. However, they generally require more computing resources and additional work to set up simulations and analyze trajectories. Students submitted caDNAno design files through the CanDo web interface without writing code or using specialized local hardware. For the structures used in this module, CanDo returned predicted equilibrium shapes and flexibility maps in about 30 minutes per structure. This turnaround gave participants time to adjust inputs and rerun simulations. One participant tested helical pitches beyond the assigned values. Another varied the stiffness parameters in the model. Both examined how their changes affected the predicted tile shape.

The module is intended for upper-level undergraduates and incoming graduate students and is organized into five 3-hour sessions. Introducing caDNAno and working through the tile design takes 3–4 hours. This includes the background needed to read the scaffold and staple layout. The simulation activity takes about 2 hours, during which students can submit and compare several conditions. Tile assembly takes about 2 hours, followed by another hour to link the tiles into ribbons. Checking the assembled structures by AFM takes about 1 hour. EtBr treatment and imaging take about 2 hours for each condition. Instructors can guide students through the full workflow, from designing a tile in caDNAno to imaging the assembled structure by AFM, or teach individual stages as stand-alone activities. The design and simulation stages require only a computer with internet access. The full sequence also provides hands-on experience in DNA assembly, solution preparation, and AFM characterization.

### Limitations and Other Considerations

This pilot study involved only three participants, limiting the generality of the educational findings. Although instructor observations and participant reports provided preliminary evidence of conceptual understanding, no structured assessment was conducted. Therefore, learning gains could not be quantitatively evaluated.

Several practical factors also affect implementation. AFM is a major constraint because it requires instrument access, operator training, and substantial imaging time. Where AFM access is limited, instructors can provide pre-acquired images or restrict imaging to selected conditions. EtBr preparation, handling, and disposal must follow approved protocols, safety data sheets, and institutional requirements.

The structural results also require careful interpretation. CanDo simulations model individual tiles at specified helical pitches, while AFM images capture assembled ribbons deposited on mica. The simulations do not account for linker junctions, surface interactions, or changes in DNA stiffness caused by intercalation. Therefore, they support the predicted relationship between helicity mismatch and deformation direction but do not directly predict the twist of the assembled ribbons.

## CONCLUSIONS

We developed and piloted a five-day laboratory module that teaches how DNA helicity governs the geometry of DNA origami assemblies. All three participants carried out the workflow themselves: they designed DNA origami tile, simulated the tile deformation at different helical pitches, assembled tiles into ribbons, treated the ribbons with EtBr, and imaged their samples using AFM. The simulations predicted that a helicity mismatch deforms individual tiles. When tiles were linked into a ribbon, these small deformations led to a global twist, with the direction of tile deformation determining the handedness of the ribbon chirality. Consistent with these predictions, all three participants observed their ribbons change from right-handed to nearly flat to left-handed conformations as the EtBr concentration increased.

Instructor observations and participant reports provided preliminary evidence that participants could relate DNA helicity, crossover placement, and structural deformation. The reports also revealed a gap in understanding the role of internal stress, identifying a clear target for future instruction. Overall, the module turns an abstract design rule that crossover placement must follow DNA helicity into a structural deformation that students can predict, produce, and observe. Future implementations with structured assessments and larger cohorts will allow learning gains to be evaluated quantitatively.

## METHODS

### Materials

Staple and linker strands were purchased from Integrated DNA Technologies (IDT). The M13mp18 scaffold was obtained from Bayou Biolabs. EtBr was purchased from Bio-Rad. All other chemicals were acquired from Sigma-Aldrich. All materials were identical to those used previously^28^.

### DNA Origami Design

The rectangular tile was designed in caDNAno^17^ on a square lattice. It consists of 32 parallel double helices, with crossovers spaced every 32 bp. Two tiles with the same scaffold routing and staple layout but distinct sequences were used (termed tiles A and B). All staple and linker sequences are identical to those reported previously^28^. Samples are referred to as monomers (single tiles), dimers (AB), and ribbons (polymerized tiles).

### CanDo Finite-Element Simulation

The caDNAno design file was submitted to the CanDo web server^22^ (https://cando-dna-origami.org) with the lattice type set to square. Default parameters were used for all mechanical inputs: axial rise, 0.34 nm; helix diameter, 2.25 nm; axial stiffness, 1100 pN; bending stiffness, 230 pN·nm^2^; torsional stiffness, 460 pN·nm^2^; nick stiffness factor, 0.01; and the coarse model. Only the helical repeat (crossover spacing) parameter was varied, with values of 10.33, 10.50, 10.67, and 10.84 bp/turn. The mismatch angle *α* for each condition was calculated as defined in the Conceptual Background. The equilibrium shape and the RMSF of each structure were obtained from the CanDo output. RMSF color maps were scaled from the minimum to the 95th-percentile value of each structure.

### Origami Assembly

Monomer tiles were prepared by mixing 10 nM M13mp18 scaffold with a 3.5-fold excess of staple strands in 1× TAE buffer containing 12.5 mM magnesium acetate (TAEM buffer). The mixture was annealed from 75 to 4 °C at 1 °C/min in a thermal cycler. Dimers were prepared by adding a linker set to an equimolar mixture of tiles A and B. Elongated ribbons were prepared by adding 50 nM of each linker strand to the 10 nM tile solution and incubating at 40 °C for 1 h.

### EtBr Treatment

Ribbons were diluted to 2 nM in imaging buffer containing 50 mM 2-(N-morpholino) ethanesulfonic acid (MES), 5 mM magnesium acetate, and 200 mM NaCl (pH 6.5). EtBr was added to final concentrations of 0, 0.25, 1, and 2 µM. EtBr was prepared, handled, and disposed of according to the safety data sheet and institutional procedures.

### AFM Imaging

For AFM, 10 µL of each sample was deposited on freshly cleaved mica and incubated for ∼10 min at room temperature in a closed Petri dish. The solution was removed with compressed air. The surface was then incubated with 10 µL of TAEM buffer for 1 min, blown dry, and rinsed with 80 µL of deionized water. Imaging was performed in air with a Dimension Icon AFM and ScanAsyst-Air probes (Bruker).

### Quantitative Analysis of Ribbon Chirality

The kink-distance data in Fig. 4c,d were taken from our previous study of the same ribbon system^28^. In that study, kink centers were marked manually in ImageJ, and the distances (d) between adjacent kink centers were measured. Positive and negative signs were assigned to right- and left-handed kinks, respectively. A total of 54 and 52 distances were measured at 0 and 2 μM EtBr, respectively, and their distributions were fitted with Gaussian functions. Kink density was calculated as the inverse of the mean signed kink-to-kink distance (1/d), and error bars indicate the standard deviation of the distances. The zero crossing of a linear fit of kink density versus EtBr concentration was used to estimate the transition between opposite ribbon conformations. These data served as an instructional example and were not generated by the participants.

### Learner Cohort, Instructional Setting, and Assessment

The module was piloted with three participants (S1–S3) who had recently earned Bachelor’s degrees in Engineering. The module was conducted over five days, with three hours of instruction and laboratory activities each day. Participants followed the workflow shown in Fig. 1a. They designed DNA origami tiles using caDNAno, performed CanDo simulations, assembled tiles and ribbons, treated ribbons with EtBr, and imaged their samples using AFM. Learning outcomes were evaluated qualitatively through participant progress reports and instructor observations during discussions. No structured pre- or post-assessment was conducted.

## ASSOCIATED CONTENT

### Data Availability Statement

The Supporting data are available on GitHub (https://github.com/choidnalab/Lab-Module-Dna-helicity-to-twist) or from the corresponding author upon reasonable request.

### Notes

The authors declare no competing financial interest.

## ACKNOWLEDGMENTS

This work was financially supported by the U.S. Department of Energy, Office of Science, Basic Energy Sciences under award no. DE-SC0020673.

